# Waves of chromatin modifications in mouse dendritic cells in response to LPS stimulation

**DOI:** 10.1101/066472

**Authors:** Alexis Vandenbon, Yutaro Kumagai, Yutaka Suzuki, Kenta Nakai

## Abstract

**Background:** The importance of transcription factors (TFs) and epigenetic modifications in the control of gene expression is widely accepted. However, causal relationships between changes in TF binding, histone modifications, and gene expression during the response to extracellular stimuli are not well understood. Here, we analyzed the ordering of these events on a genome-wide scale in dendritic cells (DCs) in response to lipopolysaccharide (LPS) stimulation.

**Results:** Using a ChIP-seq time series dataset, we found that the LPS-induced accumulation of different histone modifications follow clearly distinct patterns. Increases in H3K4me3 appear to coincide with transcriptional activation. In contrast, H3K9K14ac accumulates early after stimulation, and H3K36me3 at later time points. Integrative analysis with TF binding data revealed potential links between TF activation and dynamics in histone modifications. Especially, LPS-induced increases in H3K9K14ac and H3K4me3 were associated with binding by STAT1/2, and were severely impaired in *Stat1*^-/-^ cells.

**Conclusions:** While the timing of short-term changes of some histone modifications coincides with changes in transcriptional activity, this is not the case for others. In the latter case, dynamics in modifications more likely reflect strict regulation by stimulus-induced TFs, and their interactions with chromatin modifiers.

## Background

Epigenetic features, such as histone modifications and DNA methylation, are thought to play a crucial role in controlling the accessibility of DNA to RNA polymerases. Associations have been found between histone modifications and both long-term and short-term cellular processes, including development, heritability of cell type identity, DNA repair, and transcriptional control [1,2]. For cells of the hematopoietic lineage, cell type-defining enhancers are established during differentiation by priming with the H3K4me1 marker [3,4]. After differentiation, signals from the surrounding tissue environment or from pathogens induce changes in histone modifications reflecting the changes in activity of enhancers and promoters, including the *de novo* establishment of latent enhancers [5–9].

TFs are key regulators in the control of epigenetic changes [10,11]. During the long-term process of differentiation, closed chromatin is first bound by pioneer TFs, which results in structural changes that make it accessible to other TFs and RNA polymerase II (Pol2) [6,12]. Similarly, short-term changes in gene expression following stimulation of immune cells are regulated by TFs. This regulation is thought to involve TF binding, induction of changes in histone modifications, and recruitment of Pol2 [13–16]. However, details of the temporal ordering and causal relationships between these events remain poorly understood [17,18]. Especially, it is unclear whether certain histone modifications are a requirement for, or a result of, TF binding and transcription [19–21].

As sentinel cells of the innate immune system, DCs are well equipped for detecting the presence of pathogens. Lipopolysaccharide (LPS), a component of the cell wall of Gram negative bacteria, is recognized by DCs through the membrane-bound Toll-like receptor 4 (TLR4), resulting in the activation of two downstream signaling pathways [22]. One pathway is dependent on the adaptor protein MyD88, and leads to the activation of the TF NF-κB, which induces expression of proinflammatory cytokines. The other pathway involves the receptor protein TRIF, whose activation induces phosphorylation of the TF IRF3 by TBK1 kinase. The activated IRF3 induces expression of type I interferon, which in turn activates the JAK-STAT signaling pathway, by binding to the type I IFN receptor (IFNR) [23].

Here, we present a large-scale study of short-term changes in histone modifications in mouse DCs during the response to LPS. We focused on the timing of increases in histone modifications at promoters and enhancers, relative to the induction of transcription and to TF binding events. We observed that LPS stimulation induced increased levels of H3K9K14ac, H3K27ac, H3K4me3 and H3K36me3 at LPS-induced promoters and enhancers. Surprisingly, we observed clearly distinct patterns: accumulation of H3K9K14ac was early (between 0.5 and 2 hours after stimulation), regardless of the timing of transcriptional induction of genes. Accumulation of H3K36me3 was late, and spreads from the 3’ end of gene bodies towards the 5’ end, reaching promoters at later time points (between 8 and 24 hours). H3K4me3 accumulation was later than that of H3K9K14ac (between 1 and 4 hours), and was more correlated with transcriptional induction times. Integrated analysis with genome-wide binding data for 24 TFs revealed possible associations between increases in H3K9K14ac and H3K4me3 and binding by Rela, Irf1, and especially STAT1/2. LPS-induced accumulation of H3K9K14ac and H3K4me3 was severely impaired in *Stat1*^-/-^ cells. Together, these results suggest that stimulus-induced dynamics in a subset of histone modifications reflect the timing of activation of stimulus-dependent TFs, while others are more closely associated with transcriptional activity.

## Results

### Genome-wide Measurement of Histone Modifications at Promoter and Enhancer Regions

To elucidate the temporal ordering of stimulus-induced changes in transcription and chromatin structure, we performed chromatin immunoprecipitation experiments followed by high-throughput sequencing (ChIP-seq) for the following histone modifications in mouse DCs before and after LPS stimulation: H3K4me1, H3K4me3, H3K9K14ac, H3K9me3, H3K27ac, H3K27me3, H3K36me3, and similarly for Pol2 (Fig. S1), for ten time points (0h, 0.5h, 1h, 2h, 3h, 4h, 6h, 8h, 16h, 24h). We integrated this data with publicly available whole-genome transcription start site (TSS) data (TSS-seq) [24]. All data originated from the same cell type, treated with the same stimulus, and samples taken at the same time points. Snapshots of the data for a selection of features at four promoters are shown in Fig. S2.

Using this data collection, we defined 24,416 promoters (based on TSS-seq data and Refseq annotations) and 34,079 enhancers (using H3K4me1^high^/H3K4me3^low^ signals) (see Methods). For this genome-wide set of promoters and enhancers, we estimated the levels of histone modifications, Pol2 binding, and RNA reads over time (see Methods).

### Epigenetic Changes at Inducible Promoters and their Enhancers

Recent studies using the same cell type and stimulus showed that most changes in gene expression patterns were controlled at the transcriptional level, without widespread changes in RNA degradation rates [25,26]. We therefore defined 1,413 LPS-induced promoters based on increases in TSS-seq reads after LPS stimulation. Similarly, for both promoters and enhancers, we defined significant increases in histone modifications and Pol2 binding by comparison to pre-stimulation levels. Our analysis suggested that changes were in general rare; only 0.7 to 5.3 % of all promoters (Fig. 1A) and 0.2 to 11.0 % of all enhancers (Fig. 1B) experienced significant increases in histone modifications and Pol2 binding. However, changes were frequent at LPS-induced promoters, especially for markers of activity such as Pol2 binding, H3K4me3, H3K27ac, and H3K9K14ac, as well as for H3K36me3 (Fig. 1A). For example, while only 957 promoters (out of a total of 24,416 promoters; 3.9%) experienced significant increases in H3K9K14ac, this included 27.6% of the LPS-induced promoters (390 out of 1,413 promoters). To a lesser extent, we observed the same tendency at associated enhancers (Fig. 1B). The smaller differences at enhancers are likely to be caused by imperfect assignments of enhancers to LPS-induced promoters (i.e. we naively assigned enhancers to their most proximal promoter). Analysis of an independent ChIP-seq dataset originating from LPS-treated macrophages [6] revealed a high consistency between DCs and macrophages in LPS-induced increases in Pol2 binding, H3K27ac, and H3K4me3 at promoters and enhancers (see Supplementary Results section “Analysis of histone modification changes in LPS-treated macrophages”, and Fig. S3). The overlap in increases in H3K4me1 at enhancers was lower, though still statistically significant (p < 1e-4, based on 10,000 randomizations), possibly reflecting differences between DC- and macrophage-specific enhancers and the molecular processes that define these cell types.

**Figure 1:**
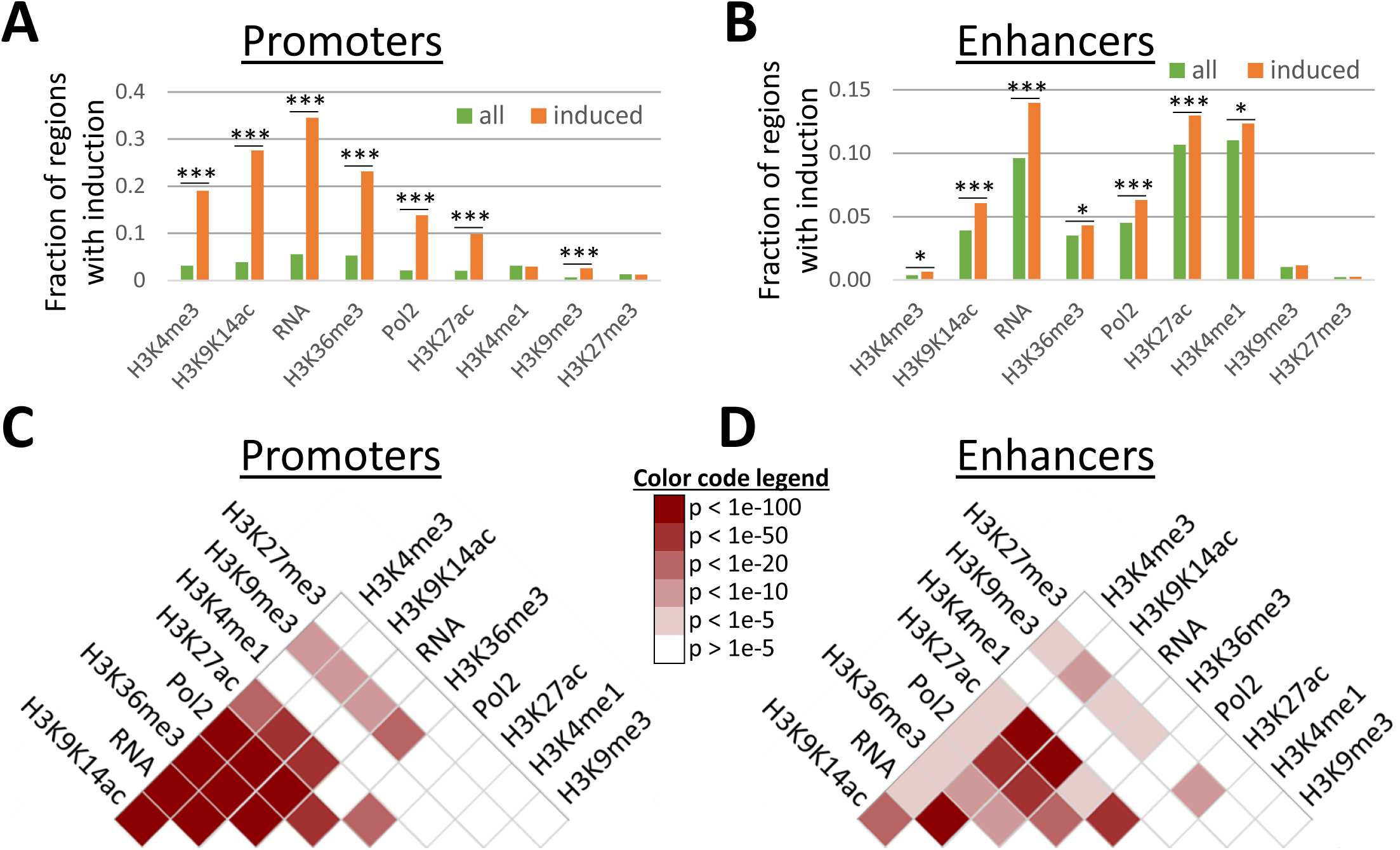
Frequencies of induction of features at LPS-induced promoters. (A) The fraction of promoters (y axis) with increases in features (x axis) are shown for the genome-wide set of promoters (green), and for the LPS-induced promoters (orange). Increases in H3K4me3, H3K9K14ac, H3K27ac, H3K36me3, RNA, and Pol2 binding are observed frequently at LPS-induced promoters. Significance of differences was estimated using Fisher’s exact test; *: p < 1e-4; **: p < 1e-6; ***: p < 1e-10. (B) Same as (A), for enhancers. (C-D) Heatmaps indicating the overlap in induction of pairs of features. Colors represent p values (-log_10_) of Fisher’s exact test. White: low overlap; Red: high overlap. Plots are shown for promoters (C), and enhancers (D).

LPS-induced promoters were less frequently associated with CpG islands (57%) than stably expressed promoters (87%, Fig. S4A) [27]. Non-CpG promoters more frequently had lower basal levels (i.e. levels at 0h, before stimulation) of activation-associated histone modifications, such as H3K27ac, H3K9K14ac, H3K4me3, and similarly lower levels of Pol2 binding and pre-stimulation gene expression (Fig. S4B). This partly explains the higher frequency of significant increases in histone modifications at LPS-induced promoters (Fig. 1A), and the higher fold-induction of genes associated with non-CpG promoters (Fig. S4C).

Previous studies have reported only limited combinatorial complexity between histone modifications, i.e. subsets of modifications are highly correlated in their occurrence [28,29]. In our data too, basal levels of activation markers at promoters and, to a lesser degree, at enhancers were highly correlated (Fig. S5). Stimulus-induced accumulations of histone modifications and Pol2 binding at promoters and enhancers further support this view: increases in H3K9K14ac, H3K4me3, H3K36me3, H3K27ac, Pol2 binding, and transcription often occurred at the same promoters (Fig. 1C). Similarly, increases in H3K9K14ac, H3K27ac, Pol2 binding, and transcription often coincided at enhancer regions (Fig. 1D). In general, activated regions experienced increases in several activation markers.

### Several Histone Modifications are Induced at a Specific Time after Stimulation

Previous studies have reported considerable dynamics in histone modifications in response to environmental stimuli (see Introduction) based on the analysis of small numbers of time points. Our dataset, however, allows the analysis of the order and timing of changes over an extended time period after stimulation. To this end, we analyzed the induction times of transcription activity, Pol2 binding, and histone modifications.

First, we inferred the transcriptional induction time of the 1,413 LPS-induced genes (see Methods and Fig. 2A). In addition, we defined a set of 772 promoters with highly stable activity over the entire time course.

**Figure 2:**
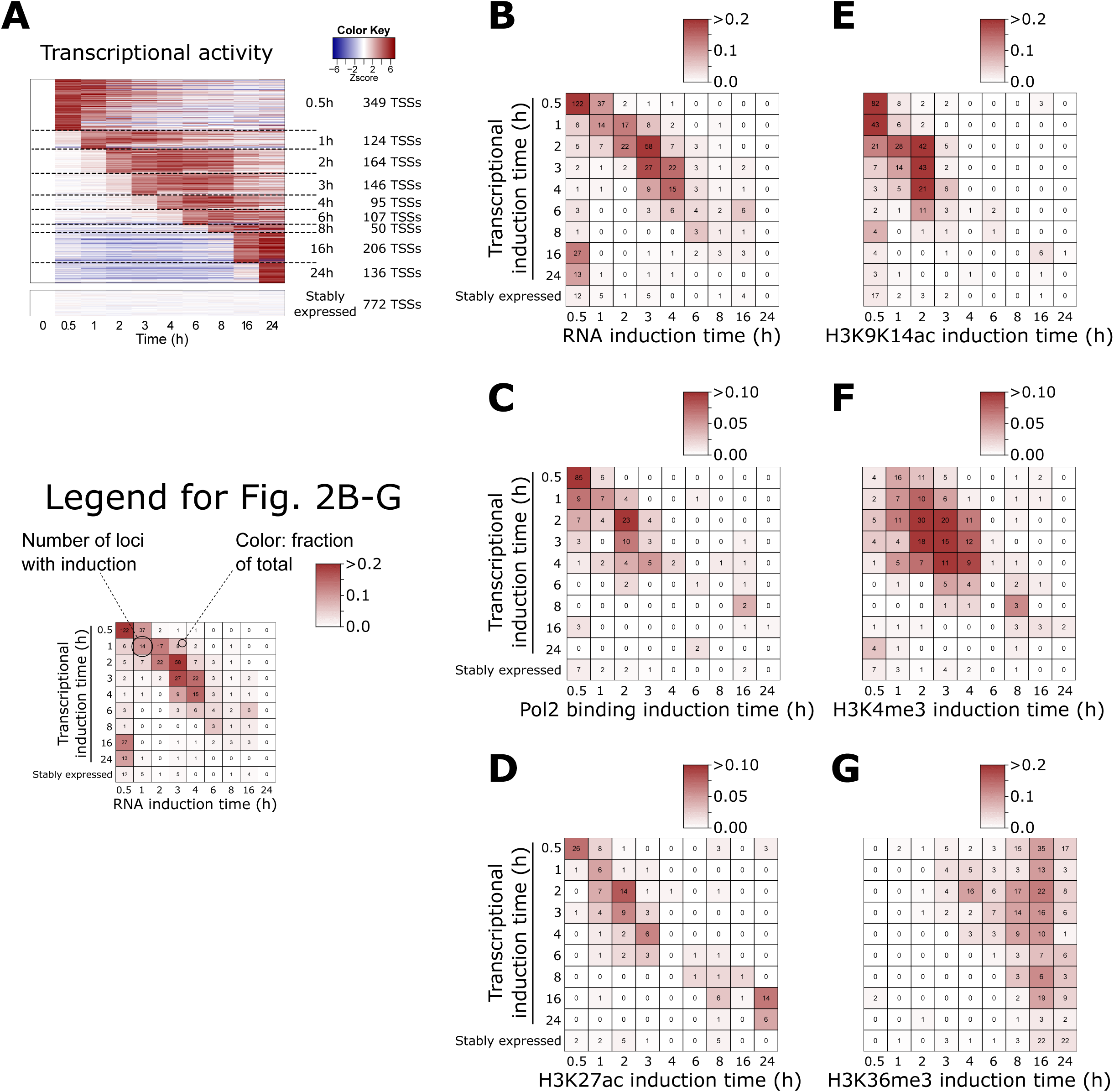
Induction times of transcription, Pol2 binding and histone modifications at promoters in function of induction of transcriptional activation times. (A) Heatmap showing the changes (white: no change; red: induction; blue: repression) in transcriptional activity of 1,413 LPS-induced promoters and stably expressed promoters, relative to time point 0h. At the right, induction times and the number of promoters induced at each time point are indicated. (B-G) Timing of increases in RNA-seq reads (B), Pol2 binding (C), H3K27ac (D), H3K9K14ac (E), H3K4me3 (F), and H3K36me3 (G) at LPS-induced promoters are shown in function of their transcription initiation times. The count of promoters with increases are indicated. Colors represent the fraction of promoters per transcriptional induction time.

As a proof of concept, using an independent time series of RNA-seq samples, we confirmed that significant increases in RNAs are seen at LPS-induced promoters in a consistent temporal order (Fig. 2B). E.g. at promoters with early induction of transcription initiation (TSS-seq) there was an early induction of mapped RNA-seq reads, while those with later induction have later induction of mapped reads. Promoters of stably expressed genes lack induction of mapped RNA reads at their promoter. Significant increases in Pol2 binding were less frequent, but followed a similar pattern (Fig. 2C).

However, the accumulation of histone modifications showed more varying patterns (Fig. 2D-G). Increases in H3K9K14ac were in general early, between 0.5h and 2h after stimulation (Fig. 2E), although promoters with early induction of transcription (0.5h, 1h, 2h) tended to have early increases in H3K9K14ac (at 0.5h). Even genes with transcriptional induction between 3-6 hours had increases in H3K9K14ac between 0.5-2 hours after stimulation. Therefore, the increases in acetylation for these promoters *preceded* induction of transcription. Significant increases later than 3 hours after stimulation were rare. In addition, increases were rare at promoters with late induction (16h, 24h) or at stably active promoters.

Increases in H3K4me3 were concentrated between 1 and 4 hours after stimulation (Fig. 2F). In contrast with H3K9K14ac, increases in H3K4me3 were rare at time point 0.5h. Accumulation of H3K4me3 was frequent at promoters with transcriptional induction between 1 and 4 hours, but – in contrast with H3K9K14ac – it was rare at immediate-early promoters.

Finally, H3K36me3 was only induced at later time points (between 8h and 24h), regardless of transcriptional induction times of promoters (Fig. 2G). In contrast with H3K9K14ac and H3K4me3, H3K36me3 is located within gene bodies and peaks towards their 3’ end (Fig. S6) [30]. Upon stimulation, H3K36me3 gradually accumulated within the gene bodies of LPS-induced genes, spreading towards the 5’ end, and reached the promoter region at the later time points in our time series (Fig. S6A). Stably expressed genes had on average high basal levels of H3K36me3, with only limited changes over time. However, interestingly, even for stably expressed genes an accumulation of H3K36me3 was observed towards their 5’ end at time points 16-24h (Fig. S6B).

Remarkably, the induction times of H3K9K14ac, H3K4me3, and H3K36me3 at promoters did not change depending on their basal levels (Fig. S7); regardless of their pre-stimulus levels, increases in H3K9K14ac were early, followed by H3K4me3, and H3K36me3 accumulation was late. This might indicate that a common mechanism is regulating these accumulations, regardless of basal levels. No differences in the accumulation times were observed between non-CpG promoters and CpG island-associated promoters (Fig. S8).

Compared to H3K9K14ac, H3K4me3 and H3K36me3, significant increases in H3K27ac appeared to be less frequent at promoters (Fig. 1A), and their timing coincided with the induction of transcription (Fig. 2D). Increases in H3K9me3, H3K27me3, and H3K4me1 were rare at promoters (Fig. S9).

The early accumulation of H3K9K14ac, followed by H3K4me3, were confirmed using an independent replicate TSS-seq data and ChIP-seq time series dataset (Fig. S10). Although accumulation of both modifications was earlier in the duplicate data than in the original time series, their relative ordering was preserved. Additional replication was performed using RT-qPCR and ChIP-qPCR measuring RNA, H3K9K14ac, H3K4me3 (see WT data in Fig. S11), and H3K36me3 (Fig. S12) at the promoters of 9 LPS-induced genes. Here too, accumulation of H3K9K14ac and H3K4me3 occurred early, with H3K9K14ac preceding H3K4me3, while accumulation of H3K36me3 occurred later. The early (between 0 and 4h) timing of LPS-induced increases in H3K27ac and H3K4me3 were further supported by the analysis of an independent ChIP-seq time series dataset (0, 4, and 24h) originating from LPS-treated macrophages [6] (see Supplementary Results section “Analysis of histone modification changes in LPS-treated macrophages”, and Fig. S13).

### Similarities and Differences between Enhancer and Promoter Induction Patterns

Interactions between enhancers and promoters could allow histone modifiers at promoters to also affect modifications at enhancer regions, and vice versa. We therefore analyzed induction of features at enhancers in function of induction of transcription at promoters (Fig. S14). In brief, we found that enhancer-associated transcripts are frequently induced at early time points (0.5 and 1h; Fig. S14A). These observations fit with those reported in a recent study showing that transcription at enhancers precedes that of promoters in cells treated with various stimuli [31]. For H3K9K14ac, the pattern at enhancers was similar to that of promoters (Fig. S14D); increases in H3K9K14ac were mainly concentrated between 0.5h and 2h after stimulation. Increases in H3K27ac at enhancers appeared to follow to some degree the order of transcription induction of nearby promoters (Fig. S14C). For other modifications, there are discrepancies between promoters and enhancers (see Fig. S14E-I).

### Correlation between LPS-induced TF Binding and Increases in Epigenetic Features

To reveal potential regulatory mechanisms underlying the epigenetic changes induced by LPS, we performed an integrative analysis of our histone modification data with TF binding data. For this we used a publicly available ChIP-seq dataset for 24 TFs with high expression in mouse DCs [32], before and after treatment with LPS (typical time points include 0h, 0.5h, 1h, and 2h, see Methods).

Initial analysis confirmed the known widespread binding of promoters by PU.1 and C/EBPβ, and to a lesser degree by IRF4, JUNB, and ATF3 [32] (Fig. S15A), and the known association between H3K4me1 and binding by PU.1 and C/EBPβ (Fig. S16A,B) [12,15]. LPS-induced promoters were frequently bound by TFs controlling the response to LPS, such as NF-κB (subunits NFKB1, REL, and RELA) and STAT family members (Fig. S15B).

Focusing on the overlap between LPS-induced TF binding at promoters and enhancers, and accumulation of epigenetic features, we found that binding of promoters by RelA, IRF1, STAT1, and STAT2 was especially associated with increases in H3K9K14ac, H3K4me3, H3K36me3, transcription, and to a lesser degree Pol2 binding and H3K27ac (Fig. 3, left; Fisher’s exact test). For example, of the 418 promoter regions that become newly bound by STAT1 after stimulation, 223 (53.3%) experience increases in H3K9K14ac (vs 3.0% of promoters not bound by STAT1; p: 8.3E-205). LPS-induced binding by the same four TFs was also strongly associated with increases in H3K9K14ac and H3K27ac at enhancers (Fig. 3, right). Combinations of these four TFs often bind to the same promoters and enhancers (Fig. S15C,D), and STAT1 functions both as a homodimer or as a heterodimer with STAT2 [33]. LPS-induced TFs, including NF-κB and STAT family members, have been shown to bind preferentially at loci that are pre-bound by PU.1, C/EBPβ, IRF4, JUNB, and ATF3 [32]. Accordingly, histone modifications were also more frequently observed at regions that were pre-bound by these five TFs (Fig. S17).

**Figure 3:**
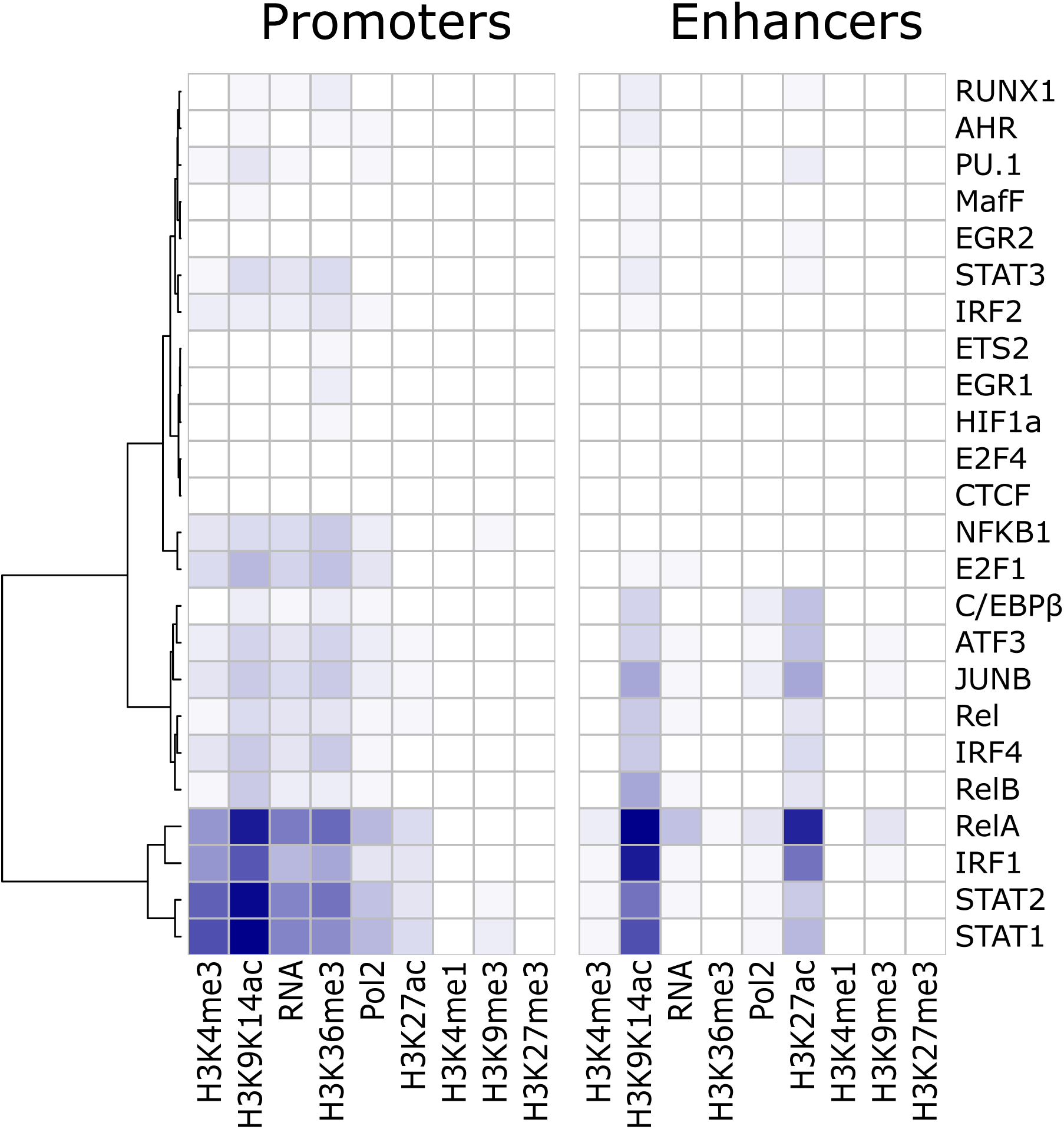
Associations between LPS-induced TF binding at promoters (left) and enhancers (right) and increases in histone modifications, Pol2 binding and transcription at the newly bound regions. Colors in the heatmap represent the degree of co-incidence (Fisher’s exact test, −log_10_ p values) between new TF binding events (rows) and increases (columns). TFs (rows) have been grouped through hierarchical clustering by similarity.

Weaker associations were found for LPS-induced binding by other NF-κB subunits (NFKB1, REL, and RELB), TFs with pervasive binding even before stimulation (C/EBPβ, ATF3, JUNB, and IRF4), and E2F1, which has been shown to be recruited by NF-ĸB through interaction with Rela [34].

Together, these results suggests a strong correlation between increases in activation marker histone modifications and LPS-induced binding by RelA, IRF1, STAT1 and STAT2.

### STAT1 and STAT2 Binding Coincides with Accumulation of H3K9K14ac, and Precedes Accumulation of H3K4me3

The relative timing of LPS-induced TF binding events and increases in histone modifications can reflect potential causal relationships. Particularly, many LPS-induced promoters show increases in H3K9K14ac between 0.5 and 2 hours after LPS stimulation (Fig. 2E), and we found a strong overlap between increases in H3K9K14ac and binding by STAT1 (Fig. 3). STAT1 is not active before stimulation, and its activity is only induced about 2 hours after LPS stimulation [35], resulting in a strong increase in STAT1-bound loci (from 56 STAT1-bound loci at 0h to 1,740 loci at 2h; Fig. S15B).

We observed a particularly strong coincidence in timing between STAT1 binding and increases in H3K9K14ac (Fig. 4A): genomic regions that become bound by STAT1 at 2h show a coinciding sharp increase in H3K9K14ac around the STAT1 binding sites. At promoters and enhancers that became bound by STAT1 at 2h the induction of H3K9K14ac was particularly frequent (Fig. 4B,C). Of the 407 promoters and 378 enhancers that become bound by STAT1 at 2 hours after stimulation, 222 (54%) and 214 (57%) have an increase in H3K9K14ac (versus only 3.0% of promoters and 3.3% of enhancers lacking STAT1 binding). These increases were especially frequent at the 2 hour time point (Fig. 4B,C).

**Figure 4:**
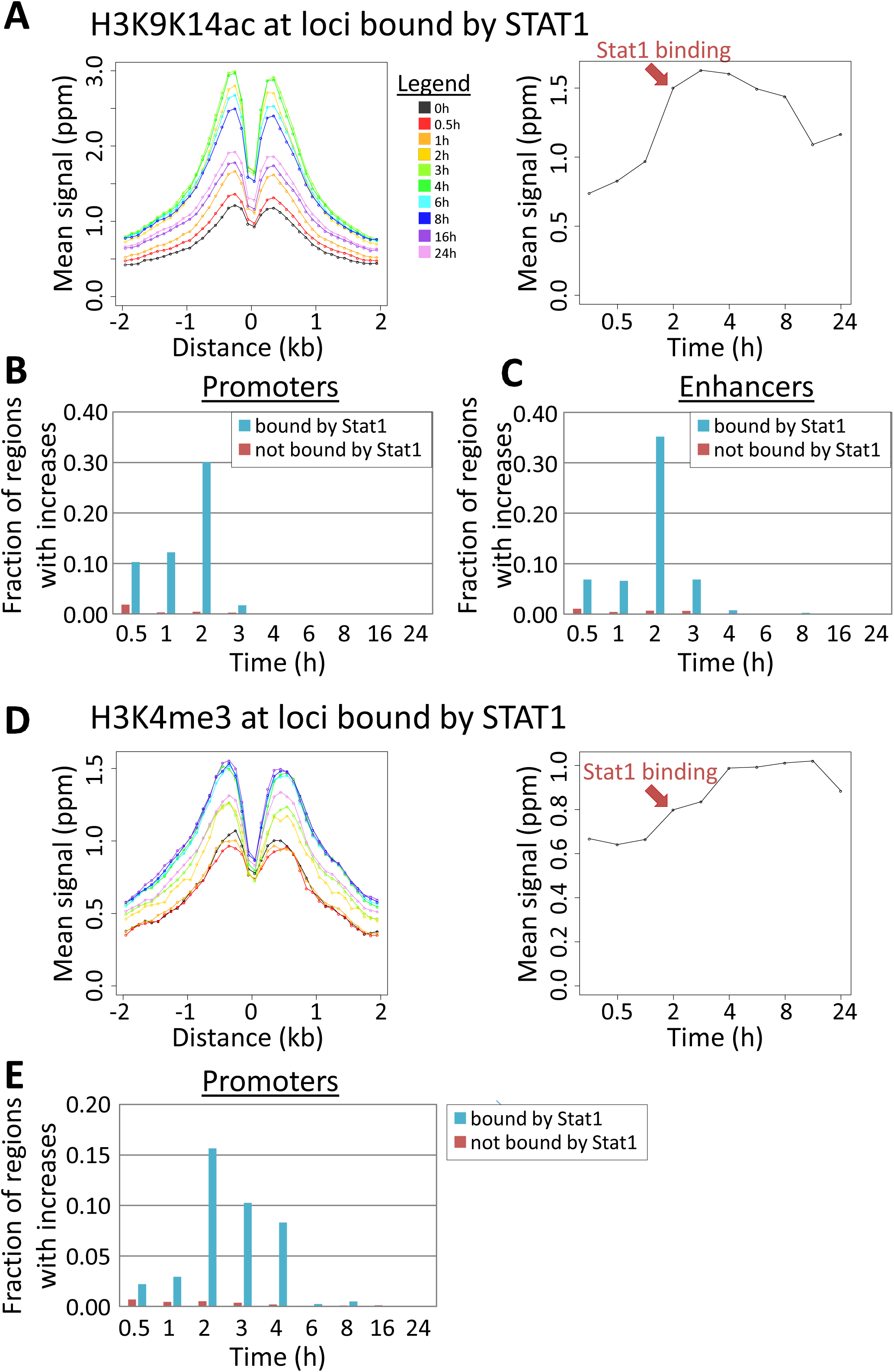
Interaction between STAT1 binding and accumulation of H3K9K14ac (A-C) and H3K4me3 (D-E). (A) For all genomic regions bound by STAT1 at 2h after LPS stimulation, mean H3K9K14ac signals are shown over time. Left: profile of mean values (y axis) over time in bins of 100 bps in function of distance (x axis) to the TF binding site. Right: mean values (y axis) summed over the region −2kb to +2kb over all bound regions, over time (x axis). The red arrow indicates the time at which these regions become bound by STAT1. (B) The fraction of promoters with increases in H3K9K14ac at each time point after stimulation (x axis). Blue: the 409 promoters bound by STAT1 at time 2h. Red: 23,964 promoters not bound by STAT1 at any time point. (C) As in (B) for 378 enhancer regions bound and 33,693 not bound by STAT1. (D) As in (A), for H3K4me3 at the genomic regions bound by STAT1 2 hours after LPS stimulation. (E) As in (B), for increases in H3K4me3 at 409 promoters bound by STAT1 and 23,964 promoters not bound by STAT1.

Similar to H3K9K14ac, we observed a general increase in H3K4me3 around STAT1 binding sites (Fig. 4D), between 2 to 4 hours after stimulation. Accordingly, only 21 STAT1-bound promoters (out of 409; 5.1%) had significant increases between 0.5 to 1 hour, but an additional 140 promoters (34%) experienced increases at the following time points (2-4 hours; Fig. 4E). As noted above, H3K4me3 was in general absent at enhancers.

Similar patterns were observed for enhancers and promoters bound by STAT2 2 hours after stimulation (Fig. S18). In contrast, regions bound by RelA (Fig. S19) and IRF1 (Fig. S20) showed increased levels of H3K27ac and to a lesser degree H3K9K14ac at earlier time points. Associations with H3K9K14ac induction after 2 hours were weak compared to STAT1/2. Average increases in H3K4me3 at RelA- and IRF1-bound regions were only modest (Fig. S19G-I and S20G-I), suggesting that the association between RelA- and IRF1-binding and H3K4me3 as seen in Fig. 3 is mostly through co-binding at STAT1/2-bound regions. Associations between histone modifications and binding by other TFs were in general weak (not shown; see Fig. 3). No changes were observed in H3K4me1 at STAT1/2-bound regions (Fig. S21A). Although there was a tendency for STAT1/2-bound loci to have increases in H3K27ac, binding seemed to slightly lag behind H3K27ac induction (Fig. S21B). Finally, although STAT1/2-bound regions tended to experience increases in H3K36me3, there was a large time lag between binding and increases in this modification (Fig. S21C). This is also true for other TFs, such as RelA and IRF1, and even PU.1 and C/EBPβ, regardless of the timing of TF binding (Fig. S16C-F).

These results suggest possible causal relationships between STAT1/2 binding and the accumulation of H3K9K14ac and H3K4me3. The specific timing of increases in these modifications might reflect the timing of activation of these TFs, resulting in the recruitment of acetyl transferases and methyl transferases to specific promoter and enhancer regions.

### LPS-induced changes in H3K9K14ac and H3K4me3 are strongly affected in *Stat1*^-/-^ cells

We decided to further investigate the role of STAT1 in controlling the changes in histone modifications. In *Trif*^-/-^ knock out (KO) cells, LPS-induced type I IFN production, activation of the JAKSTAT pathway, and activation of STAT1 and STAT2 target genes are severely impaired [23]. Using *Trif*^-/-^ and *MyD88*^-/-^ DCs, we defined a set of TRIF-dependent genes (Fig. S22A), and confirmed that they were frequently bound by STAT1/2 (Fig. S22B). We observed that promoters of TRIF-dependent and STAT1/2-bound genes frequently had LPS-induced increases in H3K9K14ac and H3K4me3 (Fig. S22C,D).

RT-qPCR and ChIP-qPCR experiments in WT, *Trif*^-/-^, *Irf3*^-/-^, and *Ifnar1*^-/-^ cells showed that a subset of TRIF-dependent and STAT1/2-bound genes (in particular *Ifit1* and *Rsad2*) showed increases in H3K9K14ac and H3K4me3 in WT, but not in KO cells (Fig. S11; see Supplementary Results section “A Subset of STAT1/2 Target Genes lack Induction of H3K9K14ac and H3K4me3 in *Trif*^-/-^, *Irf3*^-/-^, and *Ifnar1*^-/-^ cells”).

Furthermore, stimulation of WT cells using IFN-β induced expression of *Ifit1* and *Rsad2*, and accumulation of H3K9K14ac and H3K4me3 at their promoters (Fig. S23B). In this system, the activation of the IFNR signaling pathway, and of STAT1/2, is independent of TRIF. Accordingly, this IFN-β-induced accumulation of H3K9K14ac and H3K4me3 was not affected in *Trif*^-/-^ cells, further supporting a role for STAT1/2 in the control of these modifications at these genes.

Finally, we performed new ChIP-seq analysis of H3K9K14ac and H3K4me3 in WT and *Stat1*^-/-^ DCs (Fig. 5). Genomic regions that are bound by STAT1 showed a sharp increase in H3K9K14ac (Fig. 5A) and H3K4me3 (Fig. 5D) in WT cells, reproducing our observations from our first time series data (Fig. 4A,D). However, this increase was completely abrogated in *Stat1*^-/-^ cells (Fig. 5A,D). Focusing on promoter sequences, we noted 321 promoters that had increases in H3K9K14ac in WT but not in KO (Fig. 5B). These promoters were frequently bound by STAT1/2, IRF1 and NF-ĸB in WT cells (Fig. 5C). On the other hand, 184 promoters had increases in H3K9K14ac in the *Stat1*^-/-^ cells but not in WT (Fig. 5B). These promoters lack binding by STAT1/2 in WT (Fig. 5C), and the KO-specific increase in H3K9K14ac might be the result of a different set of TFs recruiting histone modifiers to these promoters, in the absence of functional STAT1. One such TF might be HIF1A, which binds a subset of these promoters but not promoters with H3K9K14ac increases in WT (Fig. 5C), and has been reported to be repressed by STAT1 [36]. Similar observations were made for H3K4me3 induction in WT and *Stat1*^-/-^ cells (Fig. 5E,F).

**Figure 5:**
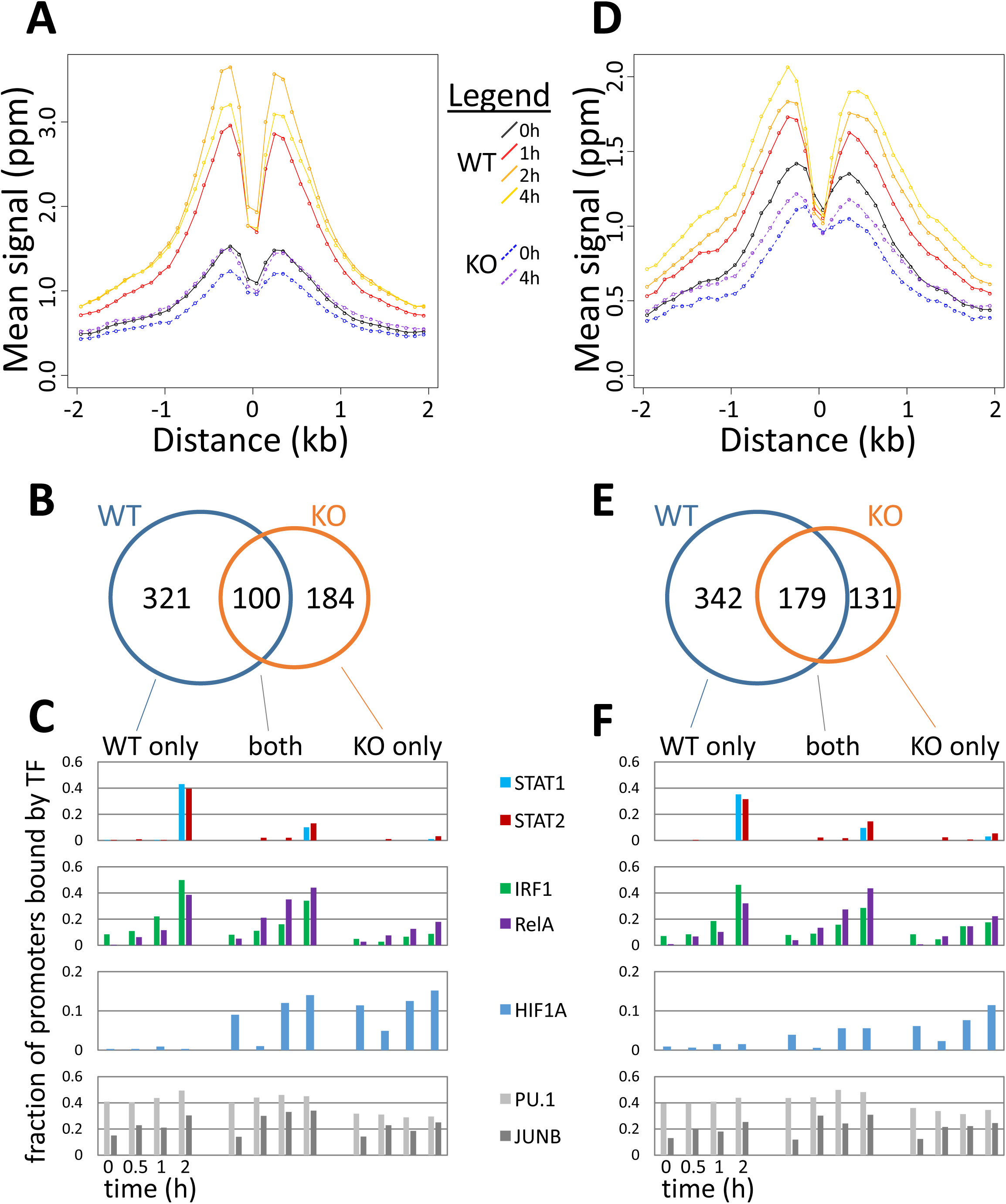
Dynamics of H3K9K14ac and H3K4me3 in *Stat1*^-/-^ cells. (A) For all genomic regions bound by STAT1 at 2h after LPS stimulation, mean H3K9K14ac signals in bins of 100 bps (y axis) are shown over time in WT and in *Stat1*^-/-^ KO cells, in function of distance (x axis) to the STAT1 binding site in WT cells. (B) A Venn diagram showing the counts of promoters with significant increases in H3K9K14ac in WT, *Stat1*^-/-^ KO, and both. (C) For promoters with increases in H3K9K14ac in WT and/or KO, the fraction bound by a selection of TFs in shown. (D-F) Same data for H3K4me3 in WT and *Stat1*^-/-^ KO cells.

## Discussion

The concept of active genes being in an open chromatin conformation was introduced several decades ago [37], but the contribution of histone modifications to the control of gene activity remains controversial [17]. In contrast, the contribution of TFs to regulating gene expression is widely recognized [38], and several studies have identified important crosstalk between TFs and histone modifiers in the regulation of the response to immune stimuli [6,7,39–43]. Nevertheless, our understanding about causal relationships between TF binding, changes in histone modifications, and changes in transcriptional activity of genes in response to stimuli is still lacking.

Analysis of the ordering of events over time can reveal insights into possible causal relationships or independence between them. Here, we have presented an integrative study of the timing and ordering of changes in histone modifications, in function of transcriptional induction in response to an immune stimulus. Our results suggest that, while the dynamics of some histone modifications are closely associated with transcriptional activity, other modifications appear to be induced at specific time frames after stimulation. For a subset of modifications (e.g. H3K9K14ac and H3K36me3) these time frames appear to be independent of the timing of induction of transcription.

In our dataset, we roughly observed three patterns of modifications. The first was early induction of H3K9K14ac, which occurs mainly in the first two hours after stimulation. A second pattern consisted of increases in H3K4me3 and H3K27ac, roughly coinciding with induction of transcription. Finally, a third pattern consisted of changes in H3K36me3, occurring only around 8-24 hours after stimulation. Although H3K4me3 is widely used as a marker for active genes, the functional role of this modification is still unclear. For example, the deletion of Set1, the only H3K4 methyltransferase in yeast, resulted in slower growth than in wild type, but otherwise appears to have only limited effects on transcription [19]. Other studies too have reported a lack of a direct effect of H3K4me3 on transcription [20,21]. Several experiments by Cano-Rodriguez and colleagues illustrate that transcription can be transiently induced in the absence of H3K4me3, and that loci-specific induction of H3K4me3 had no or limited effect on transcription [44]. Another study showed that H3K4 methyltransferase Wbp7/MLL4 controls expression of only a small fraction of genes directly [45]. In contrast, fluorescence microscopy experiments have shown that H3K27ac levels can alter Pol2 kinetics by up to 50% [21].

Since the induction of remodeling appears to occur specifically at LPS-induced genes, it is likely that histone modifiers are recruited by one or more LPS-activated TFs to specific target regions in the genome defined by the binding specificity of the TFs. In this model, primary response regulators could control immediate stimulus-induced changes in transcription and histone modifications, while the later “waves” could depend to different degrees on 1) the process of transcription itself, 2) subsequent activation of secondary regulators, and 3) the presence of other histone modifications (Fig. 6). This fits well with our observations for STAT1/2 and the induction of H3K9K14ac and H3K4me3, within specific time frames and mostly restricted to LPS-induced promoters, and the later establishment of H3K36me3. Other studies have reported associations between STAT1 binding and changes in epigenetic markers following environmental stimulation, including the activation of latent enhancers [6] and histone acetylation [41,46]. Moreover, epigenetic priming by histone acetylation through STAT1 binding to promoters and enhancers of *Tnf, Il6*, and *Il12b* has been reported, resulting in enhanced TF and Pol2 recruitment after subsequent TLR4 activation [47]. These primed regions were reported to have sustained binding by STAT1 and IRF1 and prolonged associations with CBP/p300, and constitute a stable, stimulus-induced chromatin state. The step-by-step establishment of histone modifications could reflect one way or regulating this process, with combinations of regulators deciding whether a locus will reach a stably active/poised state or whether it will return to the basal inactive state (Fig. 6). Since TFs such as STAT1 are also known to induce gene expression, one might expect the timing of increases in histone modifications to co-occur with induction of expression. However, as we described here, and as supported by the above studies, this is not necessarily the case. Gene expression is known to be regulated by combinations of TFs, and in this study too we noticed that LPS-activated TFs such as NF-κB, IRF1 and STATs often bound to the same loci (Fig. S15), which were moreover often pre-bound by several other TFs, including PU.1 and C/EBPβ. Discrepancies between timing of expression induction and accumulation of histone modifications could be caused by different requirements for combinatorial binding. This could also explain widely-reported “non-functional” TF binding, where TF binding does not seem to affect the activity of nearby genes [48]. Such “non-functional” TF binding might instead trigger changes in histone modifications that remain unnoticed and affect gene activity in more subtle ways.

**Figure 6:**
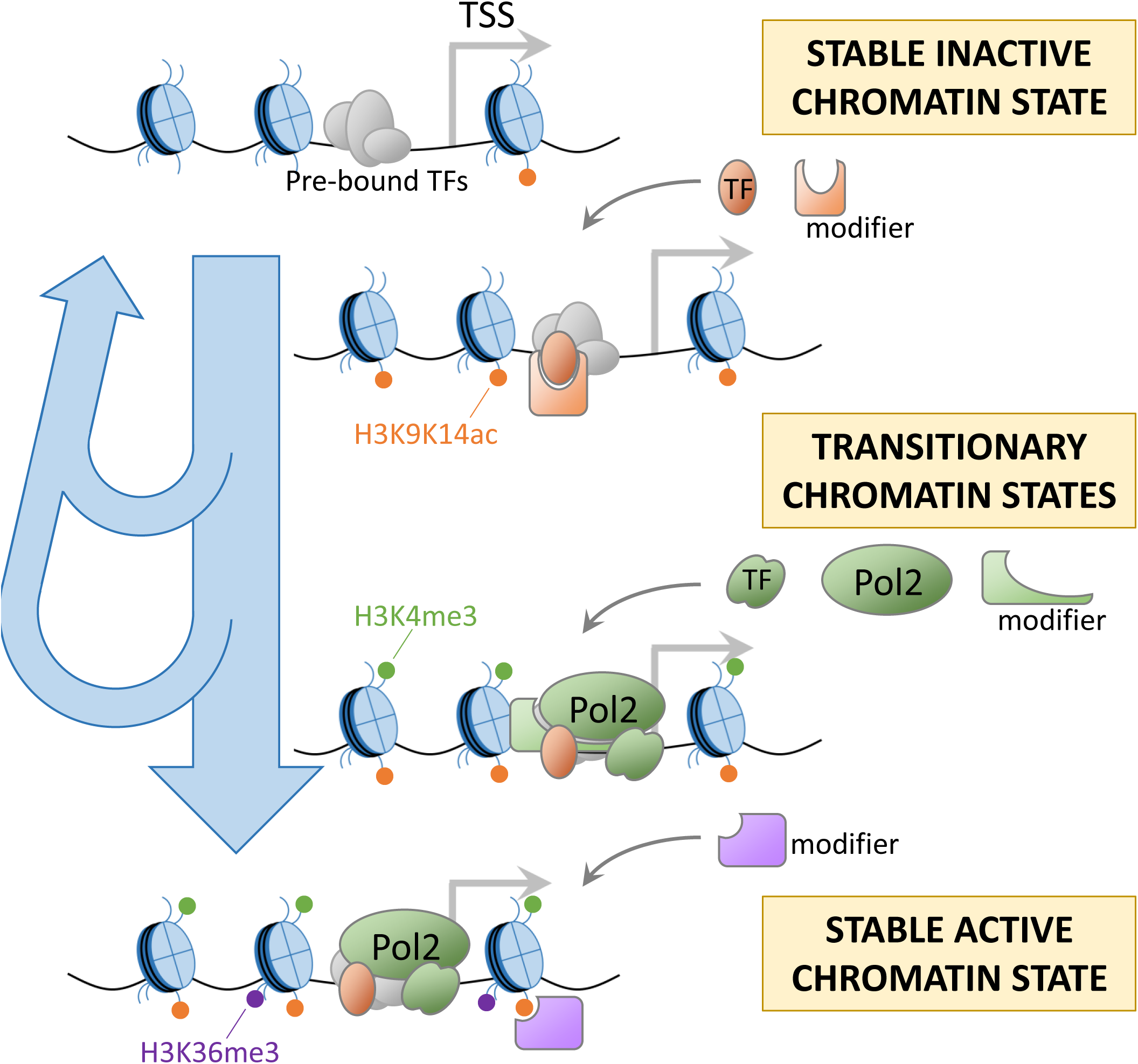
Model for the different patterns of stimulus-induced histone modifications. (i) stimulation induces the binding of primary TFs and their interacting histone modifiers (orange) at regions pre-defined by lineage-defining TFs (grey), leading to early increases in histone modifications. (ii) Secondary regulators, Pol2, and interacting histone modifiers (green) establish additional modifications at specific time points. (iii) Downstream regulators and existing histone modifications lead to further recruitment of histone modifiers (purple), establishing a stably active chromatin state.

Although many studies have compared histone modifications before and after stimulation, most lack sufficient time points and resolution to allow analysis of temporal ordering of changes. One recent study in yeast reported results that are partly similar to ours [49]: specific modifications (especially, but not only, acetylation) occur at earlier time frames during the response of yeast to diamide stress, and others at later time points. Another study in yeast showed that H3K9ac deposition appeared before the passing of the replication fork during DNA replication, while tri-methylations took more time to be established [50]. Interestingly, in these studies, typical time frames for changes in histone modifications (including H3K36me3) are less than one hour after stimulation or replication. In contrast, changes in H3K36me3 in our data appeared 8-24 hours after stimulation. Thus, time scales of stimulus-induced epigenetic changes in multicellular, higher mammalian systems might be considerably longer. Interestingly, increases in H3K36me3 around 16-24 h often coincide with a decrease in histone acetylation towards pre-stimulation levels at LPS-induced promoters. A study in yeast suggested that H3K36me3 plays a role in the activation of a histone deacetylase [51], and might therefore play a role in the return to a basal state of histone modifications and terminating the response to stimulus.

## Conclusions

Our time series ChIP-seq data and analysis present a first genome-wide view of the timing and order of accumulation of histone modifications during a stress response in mammalian immune cells. The stimulus-induced accumulation of H3K9K14ac, H3K4me3, H3K27ac, and H3K36me3 followed distinct patterns over time. Integrative analysis suggests a role for STAT1/2 in triggering increases in H3K9K14ac and H3K4me3 at stimulus-dependent promoters and enhancers. Differences in interactions between histone modifiers, TFs, and the transcriptional machinery are likely causes for the different patterns of dynamics in histone modifications.

## Material and Methods

### Reagents, cells, and mice

Bone marrow cells were prepared from C57BL/6 female mice, and were cultured in RPMI 1640 supplemented with 10 % of fetal bovine serum under the presence of murine granulocyte/monocyte colony stimulating factor (GM-CSF, purchased from Peprotech) at the concentration of 10 ng/mL. Floating cells were harvested as bone marrow-derived dendritic cells (BM-DCs) after 6 days of culture with changing medium every 2 days. The cells were stimulated with LPS (Salmonella minnessota Re595, purchased from Sigma) at the concentration of 100 ng/mL for 0, 0.5, 1, 2, 3, 4, 6, 8, 16, and 24 hours, and were subjected to RNA extraction or fixation. Murine IFN-β was purchased from Pestka Biomedical Laboratories, and was used to stimulate the cells at the concentration of 1×10^2 unit/mL. All animal experiments were approved by the Animal Care and Use Committee of the Research Institute for Microbial Diseases, Osaka University, Japan (IFReC-AP-H26-0-1-0). TRIF-, IRF3-, or IFNR-deficient mice have been described previously [52–54]. *Stat1*-deficient mouse was described previously [55].

### ChIP-seq experiments

For each time point, thirty million BM-DCs were stimulated with LPS and subjected to fixation with addition of 1/10 volume of fixation buffer (11% formaldehyde, 50 mM HEPES pH 7.3, 100 mM NaCl, 1 mM EDTA pH 8.0, 0.5 mM EGTA pH8.0). The cells were fixed for 10 minutes at room temperature, and immediately washed with PBS three times. ChIP and sequencing were performed as described (Kanai et al, DNA Res, 2011). Fifty microliter of lysate after sonication was aliquoted as “whole cell extract” (WCE) control for each IP sample. Antibodies used were Pol2 (05-623, Millipore), H3K4me3 (ab1012, Abcam), H3K9K14ac (06-599, Millipore), H3K36me3 (ab9050, Abcam), H3K9me3 (ab8898, Abcam), H3K27me3 (07-449, Milllipore), H3K4me1 (ab8895, Abcam), and H3K27ac (ab4729, Abcam).

### RNA extraction and RT-qPCR

One million BM-DCs were stimulated with LPS for indicated times and subjected to RNA extraction by using TRIzol (Invitrogen) according to manufacturer’s instruction. RNAs were reverse transcribed by using RevaTra Ace (Toyobo). The resulting cDNAs were used for qPCR by using Thunderbird SYBR master mix (Toyobo) and custom primer sets (Table S1). qPCR was performed by using LightCycler Nano (Roche).

### ChIP-qPCR

ChIP was done as above, except 4×10^6 cells were used. The resulting ChIP-DNAs were subjected to qPCR similarly to the RT-qPCR procedure, using custom primer sets (Table S2).

### Peak calling and processing of ChIP-seq data

For each histone modification and for Pol2 binding data, we aligned reads to the genome, conducted peak calling and further processing as follows.

We mapped sequenced reads of ChIP-seq IP and control (WCE) samples using Bowtie2 (version 2.0.2), using the parameter “very-sensitive”, against the mm10 version of the mouse genome [56]. Processing of alignment results, including filtering out low MAPQ alignments (MAPQ score < 30) was performed using samtools [57].

We predicted peaks for each time point using MACS (version 1.4.2) [58], using each IP sample as input and its corresponding WCE sample as control. To improve the detection of both narrow and broad peaks, peak calling was performed using default settings and also using the “nomodel” parameter with “shiftsize” set to 73. Negative control peaks were also predicted in the control sample using the IP sample as reference. Using the predicted peaks and negative control peaks, we set a threshold score corresponding to a false discovery rate (FDR) of 0.01 (number of negative control peaks vs true peaks), for each time point separately. All genomic regions with predicted peaks were collected over all 10 time points, and overlapping peak regions between time points were merged together. Moreover, we merged together peak regions separated by less than 500 bps. This gave us a collection of all genomic regions associated with a peak region in at least one sample of the time series.

In a next step, we counted the number of reads mapped to each region at each time point for both the IP samples and WCE control samples. Using these counts, we performed a read count correction, as described by Lee et al. [59]. Briefly, this method subtracts from the number of IP sample reads aligned to each peak region the expected number of non-specific reads given the number of reads aligned to the region in the corresponding WCE sample. The resulting corrected read count is an estimate of the number of IP reads in a region that would remain if no WCE reads are present [59]. This correction is necessary for the quantitative comparison of ChIP signals over time in the downstream analysis.

Finally, the corrected read counts were converted to reads per kilobase per million reads (RPKM) values (using read counts and the lengths of each region), and normalized using quantile normalization, under the assumption that their genome-wide distribution does not change substantially during each time series. The normalized RPKM values were converted to reads per million read (ppm) values.

### TSS-seq data processing and promoter definition

TSS-seq data for BM-DCs before and after stimulation with LPS was obtained from the study by Liang *et al*. [24] (DDBJ accession number DRA001234). TSS-seq data reflects transcriptional activity, but also allows for the detection of TSSs on a genome-wide scale at a 1 base resolution [60]. Mapping of TSS-seq samples was done using Bowtie2, as for ChIP-seq data. The location (5’ base) of the alignment of TSS-seq reads to the genome indicates the nucleotide at which transcription was started. In many promoters, transcription is initiated preferably at one or a few bases. Because of this particular distribution of TSS-seq reads mapped to the genome, default peak calling approaches cannot be applied. Instead, we used the following scanning window approach for defining regions with significantly high number of aligned TSS-seq reads.

The number of TSS-seq reads mapped to the genome in windows of size 1, 10, 50, 100, 500, and 1000 bases were counted in a strand-specific way, in steps of 1, 1, 5, 10, 50, and 100 bases. As a control, a large number of sequences was randomly selected from the mouse genome, and mapped using the same strategy, until an identical number of alignments as in the true data was obtained. For these random regions too, the number of reads was counted using the same scanning window approach. The distribution of actual read counts and control read counts were used to define a FDR-based threshold (FDR: 0.001) for each window size. For overlapping regions with significantly high read counts, the region with the lowest associated FDR was retained.

In order to remove potentially noisy TSSs, we removed TSSs that were located within 3’ UTRs, and TSSs located >50 kb upstream of any known gene. For remaining TSSs, we used a simple model (see Supplementary material) 1) to decide the representative TSS location in case a promoter region contained several candidate main TSSs, 2) to remove TSS-seq hits lacking typical features of promoters (e.g. presence of only TSS-seq reads in absence of histone modifications and Pol2 binding), and 3) to decide the main promoter of a gene in case there were multiple candidates. Finally, we obtained 9,964 remaining high-confidence TSSs, each assigned to 1 single Refseq gene.

These TSS-seq-based TSSs were supplemented with 14,453 non-overlapping Refseq-based TSSs for all Refseq genes which did not have an assigned high-confidence TSS-seq-based TSS. Most of the genes associated with these TSSs had lower expression in our RNA-seq data (mostly RPKM is 0 or < 1; not shown). Together, TSS-seq-based TSSs and Refseq-based TSSs resulted in a total of 24,416 promoter regions.

CpG-associated promoters were defined as those having a predicted CpG island (from the UCSC Genome Browser Database) in the region −1kb to +1kb surrounding the TSS [61]. Other promoters were considered to be non-CpG promoters.

### Definition of enhancers

Enhancers were defined based on the signals of H3K4me1 and H3K4me3. First, we collected all genomic regions with significantly high levels of H3K4me1 (see section “Peak calling and processing of ChIP-seq data”) in at least one of the ten time points. Regions located proximally (<2kb distance) to promoter regions and exons were removed, because they are likely to be weak H3K4me1 peaks observed around promoters, as were H3K4me1-positive regions of excessively large size (>10kb). Finally, we removed regions with H3K4me1 < H3K4me3 * 5, resulting in 34,072 remaining enhancers.

Enhancers were naively assigned to the nearest promoter (TSS-seq based or Refseq-based) that was < 150kb separated from it (center-to-center). For 30,448 enhancers (89%) a promoter could be assigned.

### Public ChIP-seq data for TFs

Genome-wide binding data (ChIP-seq) is available for mouse DCs before and after stimulation with LPS, for a set of 24 TFs with a known role of importance and/or high expression in DCs [32] (GEO accession number GSE36104). TFs (or TF subunits) included in this dataset are Ahr, ATF3, C/EBPβ, CTCF, E2F1, E2F4, EGR1, EGR2, ETS2, HIF1a, IRF1, IRF2, IRF4, JUNB, MafF, NFKB1, PU.1, Rel, RelA, RelB, RUNX1, STAT1, STAT2, and STAT3. Typically time points in this data are 0h, 0.5h, 1h, and 2h following LPS stimulation (some TFs lack one or more time points). We used the ChIP-seq-based peak scores and score threshold as provided by the original study as an indicator of significant TF binding.

Promoters (region −1kb to +1kb around TSS) and enhancers (entire enhancer region or region −1kb to +1kb around the enhancer center for enhancers < 2 kb in size) were considered to be bound by a TF if they overlapped a ChIP-seq peak with a significantly high peak score. New binding events by a TF at a region were defined as time points with a significantly high score where all previous time points lacked significant binding.

### Definition of induction of histone modifications and Pol2 binding

In order to analyze induction times of increases in histone modifications and Pol2 binding, we defined the induction time of a feature as the first time point at which a significant increase was observed compared to its original basal levels (at 0h). Significant increases were defined using an approach similar to methods such as by DESeq and voom [62,63], which evaluate changes between samples taking into account the expected variance or dispersion in read counts in function of mean read counts. This approach is necessary because regions with low read counts typically experience high fold-changes because of statistical noise in the data. Here we modified this approach to be applicable to our data (10 time points without replicates; ppm values per promoter/enhancer region).

The values of all histone modifications, Pol2, RNA-seq, TSS-seq reads (ppms, for each time point) were collected for all promoters (region −1kb to +1kb) and enhancers (entire enhancer region or region −1kb to +1kb around the enhancer center for enhancers < 2 kb in size). For each feature (all histone modifications and Pol2 binding), we calculated the median and standard deviation in ppm values for each region, over the 10 time points. Dispersion was defined as follow:

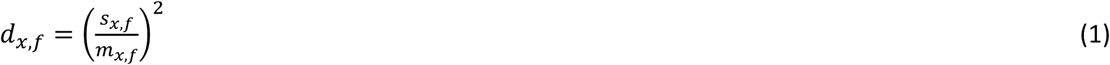

where *d_x,f_*, *s_x,f_*, and *m_x,f_* represent the dispersion, standard deviation, and median of feature *f* in region *x* over the 10 time points of the time series. Fitting a second order polynomial function on the log(*d_x,f_*) as a function of log(*m_x,f_*) for all promoter and enhancer regions, we obtained expected dispersion values in function of median ppm value (see for example Fig. S24 for H3K9K14ac). From fitted dispersion values, fitted standard deviation values *s_x,f,fitted_* were calculated (see Eq. 1), and 0h-based Z-scores were calculated as follows:

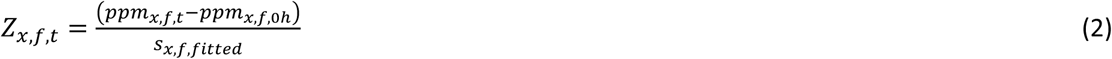

where *Z_x,f,t_* is the Z-score of feature *f* in region *x* at time point *t*, and *ppm_x,f,t_* is the ppm value of feature *f* in region *x* at time point *t*. A region *x* was defined to have a significant induction of feature *f* if there was at least one time point *t* where *Z_x,f,t_* ≥4. To further exclude low-signal regions we added this additional threshold: the region should have a ppm value ≥ the 25 percentile of non-0 values in at least 1 time point. For the regions with a significant induction, the induction time was defined as the first time point *t* where *Z_x,f,t_* ≥2. We used a similar approach to define LPS-induced promoters using TSS-seq data (see below).

For the analysis of induction times of H3K9K14ac, H3K4me3, and H3K36me3 at enhancers in function of their pre-stimulation basal levels (Fig. S7), we divided promoters into three classes according to their basal levels of each modifications as follows: Promoters lacking a modifications altogether (0 tag reads after correction described above) were considered as one class (“absent”). The remaining promoters were sorted according to their basal level of the modification, and were divided into two classes (“low basal level”, and “high basal level”) containing the same number of promoters.

### Definition of LPS-induced promoters, unchanged promoters

LPS-induced promoters were defined using TSS-seq ppm values. LPS-induced promoters should have *Z_x,TSS−seq,t_* ≥4 for at least 1 time point and have TSS-seq ppm ≥1 at at least 1 time point. Only TSS-seq reads aligned in the sense orientation were considered for this (e.g. they should fit the orientation of the associated gene). For each of the thus obtained 1,413 LPS-induced promoters, the transcription induction time was defined as the first time point for which *Z_x,TSS−seq,t_* ≥2 was observed. Unchanged promoters were defined as those promoters having absolute values of *Z_x,TSS−seq,t_* < 1 for all time points, leading to 772 promoters.

### RNA-seq data processing for wild type, *Trif*^-/-^ and *Myd88*^-/-^ cells

RNA-seq data for mouse BM-DCs treated with LPS were obtained from the study by Patil *et al.* [64] (SRA accession number DRA001131). This data includes time series data for WT, as well as *Trif*^-/-^ mice and *Myd88*^-/-^ mice.

Mapping of RNA-seq data was performed using TopHat (version 2.0.6) and Bowtie2 (version 2.0.2) [56,65]. Mapped reads were converted to RPKM values [66] using gene annotation data provided by TopHat. RNA-seq data obtained from the *Myd88*^-/-^ and *Trif*^-/-^ mice was processed in the same way. RPKM values were subjected to quantile normalization over all 10 time points.

For genes corresponding to the LPS-induced promoters, the maximum fold-induction was calculated in the WT RNA-seq data. The same was done in the *Trif*^-/-^ RNA-seq data, and in the *Myd88*^-/-^ RNA-seq data. TRIF-dependent genes were defined as genes for which the fold-induction was more than 5 times lower in the *Trif*^-/-^ data than in WT, leading to 141 TRIF-dependent genes (see Fig. S22A). Similarly, 66 MyD88-dependent genes (not shown) were defined as having more than 5 times lower induction in the *Myd88*^-/-^ than in WT.

### Duplicate ChIP-seq data and *Stat1*^-/-^ data

We generated an independent duplicate time series for dendritic cells (DCs) treated with LPS (0, 1, 2, and 4 hours), including TSS-seq, and ChIP-seq (H3K9K14ac and H3K4me3) data as described above. Data was processed in the same way as the original time series dataset (see Methods), and the induction times of H3K9K14ac and H3K4me3 at LPS-induced promoters was estimated. To facilitate the comparison between the duplicate data and the original (longer) times series, we also re-analyzed the original data using only time points 0, 1, 2, and 4 hours (the same time points as the duplicate data). Stat1 KO DCs were treated with LPS for 0 or 4 hours along with wild-type DCs, and ChIP-seq (H3K9K14ac and H3K4me3) data were obtained as described above.

### Fisher’s exact test

We used Fisher’s exact test to evaluate the significance of differences between induced and non-induced promoters and enhancers (Fig. 1A,B), the significance of associations between changes of pairs of features (Fig. 1C,D), and the association between TF binding and increases in histone modifications, Pol2 binding and transcription (Fig. 3 and Fig. S17).

## List of abbreviations

BM-DC: bone marrow-derived dendritic cells
DC: dendritic cell
FDR: false discovery rate
GM-CSF: granulocyte/monocyte colony stimulating factor IFN interferon
IFNR: interferon receptor
KO: knock out
LPS: lipopolysaccharide
Pol2: RNA polymerase II ppm reads per million reads
RPKM: reads per kilobase per million reads
TF: transcription factor
TLR: Toll-like receptor
TSS: transcription start site
WCE: whole cell extract
WT: wild type

## Declarations

### Ethics approval and consent to participate

Not applicable.

### Consent for publication

Not applicable.

### Availability of data and materials

The ChIP-seq datasets generated and analyzed during the current study are available in the DDBJ repository, accession numbers DRA004881 and DRA006555.

## Authors’ contributions

AV, YK, YS, and KN designed the project. YK and YS conducted ChIP-seq experiments, and YK additional experiments. AV and YK performed data analysis. All authors contributed to the interpretation of the data. AV and YK wrote the manuscript. All authors read and approved the final manuscript.

## Acknowledgements

We thank the members of the Quantitative Immunology Research Unit for helpful discussions and advice, A. Yoshimura, E. Kurumatani, Y. Kimura, A. Yamashita, K. Imamura, K. Abe and T. Horiuchi for technical assistance and M. Ogawa for secretarial assistance. Computational time was provided by the computer cluster of the IFReC Laboratory of Systems Immunology and in part by the NIG supercomputer at ROIS National Institute of Genetics.

## References

1. Henikoff S. Nucleosome destabilization in the epigenetic regulation of gene expression. Nat Rev Genet. 2008;9:15–26.

2. Greer EL, Shi Y. Histone methylation: a dynamic mark in health, disease and inheritance. Nat. Rev. Genet. 2012;13:343–57.

3. Mercer EM, Lin YC, Benner C, Jhunjhunwala S, Dutkowski J, Flores M, et al. Multilineage priming of enhancer repertoires precedes commitment to the B and myeloid cell lineages in hematopoietic progenitors. Immunity. 2011;35:413–25.

4. Winter DR, Amit I. The role of chromatin dynamics in immune cell development. Immunol. Rev. 2014;261:9–22.

5. Creyghton MP, Cheng AW, Welstead GG, Kooistra T, Carey BW, Steine EJ, et al. Histone H3K27ac separates active from poised enhancers and predicts developmental state. Proc. Natl. Acad. Sci. U. S. A. 2010;107:21931–6.

6. Ostuni R, Piccolo V, Barozzi I, Polletti S, Termanini A, Bonifacio S, et al. Latent enhancers activated by stimulation in differentiated cells. Cell. 2013;152:157–71.

7. Kaikkonen MU, Spann NJ, Heinz S, Romanoski CE, Allison K a, Stender JD, et al. Remodeling of the enhancer landscape during macrophage activation is coupled to enhancer transcription. Mol. Cell. 2013;51:310–25.

8. Lavin Y, Winter D, Blecher-Gonen R, David E, Keren-Shaul H, Merad M, et al. Tissue-Resident Macrophage Enhancer Landscapes Are Shaped by the Local Microenvironment. Cell. 2014;159:1312–26.

9. Gosselin D, Link VM, Romanoski CE, Fonseca GJ, Eichenfield DZ, Spann NJ, et al. Environment Drives Selection and Function of Enhancers Controlling Tissue-Specific Macrophage Identities. Cell. 2014;159:1327–40.

10. Voss TC, Hager GL. Dynamic regulation of transcriptional states by chromatin and transcription factors. Nat. Rev. Genet. 2014;15:69–81.

11. Álvarez-Errico D, Vento-Tormo R, Sieweke M, Ballestar E. Epigenetic control of myeloid cell differentiation, identity and function. Nat. Rev. Immunol. 2014;15:7–17.

12. Heinz S, Benner C, Spann N, Bertolino E, Lin YC, Laslo P, et al. Simple combinations of lineage-determining transcription factors prime cis-regulatory elements required for macrophage and B cell identities. Mol. Cell. 2010;38:576–89.

13. Foster SL, Hargreaves DC, Medzhitov R. Gene-specific control of inflammation by TLR-induced chromatin modifications. Nature. 2007;447:972–8.

14. Smale ST, Tarakhovsky A, Natoli G. Chromatin contributions to the regulation of innate immunity. Annu. Rev. Immunol. 2014;32:489–511.

15. Ghisletti S, Barozzi I, Mietton F, Polletti S, De Santa F, Venturini E, et al. Identification and characterization of enhancers controlling the inflammatory gene expression program in macrophages. Immunity. 2010;32:317–28.

16. Natoli G. Control of NF- k B-dependent Transcriptional Responses by Chromatin Organization. Cold Spring Harb Perspect Biol. 2009;1:1–11.

17. Henikoff S, Shilatifard A. Histone modification: Cause or cog? Trends Genet. 2011;27:389–96.

18. Ivashkiv LB, Park SHO. Epigenetic Regulation of Myeloid Cells. Microbiol Spectr. 2016;4.

19. Miller T, Krogan NJ, Dover J, Tempst P, Johnston M, Greenblatt JF, et al. COMPASS: A complex of proteins associated with a trithorax-related SET domain protein. Proc. Natl. Acad. Sci. U. S. A. 2001;98:12902–7.

20. Pavri R, Zhu B, Li G, Trojer P, Mandal S, Shilatifard A, et al. Histone H2B Monoubiquitination Functions Cooperatively with FACT to Regulate Elongation by RNA Polymerase II. Cell. 2006;125:703–17.

21. Stasevich TJ, Hayashi-Takanaka Y, Sato Y, Maehara K, Ohkawa Y, Sakata-Sogawa K, et al. Regulation of RNA polymerase II activation by histone acetylation in single living cells. Nature. 2014;516:272–5.

22. Kawai T, Akira S. The role of pattern-recognition receptors in innate immunity: update on Toll-like receptors. Nat. Immunol. 2010;11:373–84.

23. Hoshino K, Kaisho T, Iwabe T, Takeuchi O, Akira S. Differential involvement of IFN- in Toll-like receptor-stimulated dendritic cell activation. Int. Immunol. 2002;14:1225–31.

24. Liang K, Suzuki Y, Kumagai Y, Nakai K. Analysis of changes in transcription start site distribution by a classification approach. Gene. 2014;537:29–40.

25. Rabani M, Raychowdhury R, Jovanovic M, Rooney M, Stumpo DJ, Pauli A. Resource High-Resolution Sequencing and Modeling Identifies Distinct Dynamic RNA Regulatory Strategies. Cell. 2014;159:1698–710.

26. Kumagai Y, Vandenbon A, Teraguchi S, Akira S, Suzuki Y. Genome-wide map of RNA degradation kinetics patterns in dendritic cells after LPS stimulation facilitates identification of primary sequence and secondary structure motifs in mRNAs. BMC Genomics. 2016;17:1032.

27. Illingworth RS, Bird AP. CpG islands--’a rough guide’. FEBS Lett. 2009;583:1713–20.

28. Schübeler D, MacAlpine DM, Scalzo D, Wirbelauer C, Kooperberg C, Van Leeuwen F, et al. The histone modification pattern of active genes revealed through genome-wide chromatin analysis of a higher eukaryote. Genes Dev. 2004;18:1263–71.

29. Ernst J, Kheradpour P, Mikkelsen TS, Shoresh N, Ward LD, Epstein CB, et al. Mapping and analysis of chromatin state dynamics in nine human cell types. Nature. 2011;473:43–9.

30. Barski A, Cuddapah S, Cui K, Roh T-Y, Schones DE, Wang Z, et al. High-resolution profiling of histone methylations in the human genome. Cell. 2007;129:823–37.

31. Arner E, Daub CO, Vitting-Seerup K, Andersson R, Lilje B, Drabløs F, et al. Transcribed enhancers lead waves of coordinated transcription in transitioning mammalian cells. Science. 2015;347:1010–5.

32. Garber M, Yosef N, Goren A, Raychowdhury R, Thielke A, Guttman M, et al. A High-Throughput Chromatin Immunoprecipitation Approach Reveals Principles of Dynamic Gene Regulation in Mammals. Mol. Cell. 2012;47:810–22.

33. Ramana C V, Chatterjee-Kishore M, Nguyen H, Stark GR. Complex roles of Stat1 in regulating gene expression. Oncogene. 2000;19:2619–27.

34. Lim CA, Yao F, Wong JJY, George J, Xu H, Chiu KP, et al. Genome-wide Mapping of RELA(p65) Binding Identifies E2F1 as a Transcriptional Activator Recruited by NF-κB upon TLR4 Activation. Mol. Cell. 2007;27:622–35.

35. Toshchakov V, Jones BW, Perera P-Y, Thomas K, Cody MJ, Zhang S, et al. TLR4, but not TLR2, mediates IFN-beta-induced STAT1alpha/beta-dependent gene expression in macrophages. Nat. Immunol. 2002;3:392–8.

36. Hiroi M, Mori K, Sakaeda Y, Shimada J, Ohmori Y. STAT1 represses hypoxia-inducible factor-1-mediated transcription. Biochem. Biophys. Res. Commun. 2009;387:806–10.

37. Weintraub H, Groudine M. Chromosomal Subunits in Active Genes Have an Altered Conformation. Science. 1976;193:848–56.

38. Lenhard B, Sandelin A, Carninci P. Metazoan promoters: emerging characteristics and insights into transcriptional regulation. Nat. Rev. Genet. 2012;13:233–45.

39. Kayama H, Ramirez-Carrozzi VR, Yamamoto M, Mizutani T, Kuwata H, Iba H, et al. Class-specific regulation of pro-inflammatory genes by MyD88 pathways and IκBζ. J. Biol. Chem. 2008;283:12468–77.

40. Tartey S, Matsushita K, Vandenbon A, Ori D, Imamura T, Mino T, et al. Akirin 2 is critical for inducing inflammatory genes by bridging IκB-ζ and the SWI / SNF complex. EMBO J. 2014;33:2332–48.

41. Chen X, Barozzi I, Termanini A, Prosperini E, Recchiuti A, Dalli J, et al. Requirement for the histone deacetylase Hdac3 for the inflammatory gene expression program in macrophages. Proc. Natl. Acad. Sci. U. S. A. 2012;109:2865–74.

42. Zhu Y, van Essen D, Saccani S. Cell-Type-Specific Control of Enhancer Activity by H3K9 Trimethylation. Mol. Cell. 2012;46:408–23.

43. Stender JD, Pascual G, Liu W, Kaikkonen MU, Do K, Spann NJ, et al. Control of Proinflammatory Gene Programs by Regulated Trimethylation and Demethylation of Histone H4K20. Mol. Cell. 2012;1:1–11.

44. Cano-Rodriguez D, Gjaltema RAF, Jilderda LJ, Jellema P, Dokter-Fokkens J, Ruiters MHJ, et al. Writing of H3K4Me3 overcomes epigenetic silencing in a sustained but context-dependent manner. Nat. Commun. 2016;7:12284.

45. Austenaa L, Barozzi I, Chronowska A, Termanini A, Ostuni R, Prosperini E, et al. The Histone Methyltransferase Wbp7 Controls Macrophage Function through GPI Glycolipid Anchor Synthesis. Immunity. 2012;36:572–85.

46. Vahedi G, Takahashi H, Nakayamada S, Sun H, Sartorelli V, Kanno Y, et al. STATs Shape the Active Enhancer Landscape of T Cell Populations. Cell. 2012;151:981–93.

47. Qiao Y, Giannopoulou EG, Chan CH, Park S-H, Gong S, Chen J, et al. Synergistic activation of inflammatory cytokine genes by interferon-γ-induced chromatin remodeling and toll-like receptor signaling. Immunity. 2013;39:454–69.

48. MacQuarrie KL, Fong AP, Morse RH, Tapscott SJ. Genome-wide transcription factor binding: beyond direct target regulation. Trends Genet. 2011;27:141–8.

49. Weiner A, Hsieh T-HS, Appleboim A, Chen H V, Rahat A, Amit I, et al. High-Resolution Chromatin Dynamics during a Yeast Stress Response. Mol. Cell. 2015;58:371–86.

50. Bar-ziv R, Voichek Y, Barkai N. Chromatin dynamics during DNA replication. Genome Res. 2016;26:1245–56.

51. Drouin S, Larame´e L, Jacques P-E, Forest A, Bergeron M, Robert F. DSIF and RNA polymerase II CTD phosphorylation coordinate the recruitment of Rpd3S to actively transcribed genes. PLoS Genet. 2010;6:1–12.

52. Yamamoto M, Sato S, Hemmi H. Role of Adaptor TRIF in the MyD88-Independent Toll-Like Receptor Signaling Pathway. Science. 2003;301:640–3.

53. Saitoh T, Satoh T, Yamamoto N, Uematsu S, Takeuchi O, Kawai T, et al. Antiviral protein viperin promotes toll-like receptor 7- and toll-like receptor 9-mediated type i interferon production in plasmacytoid dendritic cells. Immunity. 2011;34:352–63.

54. Hemmi H, Kaisho T, Takeda K, Akira S. The Roles of Toll-Like Receptor 9, MyD88, and DNA-Dependent Protein Kinase Catalytic Subunit in the Effects of Two Distinct CpG DNAs on Dendritic Cell Subsets. J. Immunol. 2003;170:3059–64.

55. Masuda K, Kimura A, Hanieh H, Nguyen NT, Nakahama T, Chinen I, et al. Aryl hydrocarbon receptor negatively regulates LPS-induced IL-6 production through suppression of histamine production in macrophages. Int. Immunol. 2011;23:637–45.

56. Langmead B, Salzberg SL. Fast gapped-read alignment with Bowtie 2. Nat. Methods. 2012;9:357–9.

57. Li H, Handsaker B, Wysoker A, Fennell T, Ruan J, Homer N, et al. The Sequence Alignment/Map format and SAMtools. Bioinformatics. 2009;25:2078–9.

58. Feng J, Liu T, Qin B, Zhang Y, Liu XS. Identifying ChIP-seq enrichment using MACS. Nat. Protoc. 2012;7:1728–40.

59. Lee B, Bhinge AA, Battenhouse A, Song L, Zhang Z, Grasfeder LL, et al. Cell-type specific and combinatorial usage of diverse transcription factors revealed by genome-wide binding studies in multiple human cells. Genome Res. 2012;22:9–24.

60. Tsuchihara K, Suzuki Y, Wakaguri H, Irie T, Tanimoto K, Hashimoto S, et al. Massive transcriptional start site analysis of human genes in hypoxia cells. Nucleic Acids Res. 2009;37:2249–63.

61. Meyer LR, Zweig AS, Hinrichs AS, Karolchik D, Kuhn RM, Wong M, et al. The UCSC Genome Browser database: extensions and updates 2013. Nucleic Acids Res. 2013;41:D64–9.

62. Anders S, Huber W. Differential expression analysis for sequence count data. Genome Biol. 2010;11:R106.

63. Law CW, Chen Y, Shi W, Smyth GK. voom: precision weights unlock linear model analysis tools for RNA-seq read counts. 2014;1–17.

64. Patil A, Kumagai Y, Liang K-C, Suzuki Y, Nakai K. Linking transcriptional changes over time in stimulated dendritic cells to identify gene networks activated during the innate immune response. PLoS Comput. Biol. 2013;9:e1003323.

65. Kim D, Pertea G, Trapnell C, Pimentel H, Kelley R, Salzberg SL. TopHat2: accurate alignment of transcriptomes in the presence of insertions, deletions and gene fusions. Genome Biol. 2013;14:R36.

66. Mortazavi A, Williams B a, McCue K, Schaeffer L, Wold B. Mapping and quantifying mammalian transcriptomes by RNA-Seq. Nat. Methods. 2008;5:621–8.

